# Tissue-specific mucociliary regulatory programs in the healthy human endocervix and ectocervix

**DOI:** 10.64898/2026.11.19.767467

**Authors:** Alliya Parvez, Olha Kholod, Larkin Lucy, Rachel Looney, Tammy C. T. Lan, Leo L. Han, Brittany A. Goods

**Author notes:** CO-CORRESPONDENCE: Brittany A. Goods, PhD School of BioSciences, The University of Melbourne 30 Flemmington Road, Parkville, VIC, 3010, Leo L. Han, M.D., Ph.D., Department of Obstetrics and Gynecology Oregon Health & Science University 3181 SW Sam Jackson Park Road Portland, OR, 97239. Equal Contribution (Alliya Parvez, Olha Kholod).

## Abstract

The cervix is a dynamic mucosal barrier in the female reproductive tract that balances inflammatory and anti-inflammatory functions to regulate physiologically critical functions, including pregnancy, sperm transport, fertility, and protection of the upper reproductive tract from infection. Despite its central role in reproduction, the cellular and transcriptional organization of the healthy human cervix remains incompletely characterized, including how clinical sampling methods influence the epithelial populations recovered and how cervical mucociliary programs compare with those of pulmonary epithelium. We performed single-nucleus RNA sequencing (snRNA-seq) on paired Tischler punch biopsy and endocervical curettage (ECC) samples from four healthy, exogenous hormone-naive donors to generate a dedicated transcriptomic map of the healthy human cervix followed by an integrative computational analysis framework combining differential expression, functional enrichment, cell–cell communication, cross-tissue mapping, and regulon activity inference. This approach identified ten cell types spanning epithelial, stromal, vascular, and immune cell populations and found that sampling method strongly influenced cellular composition and transcriptional state as Tischler punch biopsy preferentially recovered basal and suprabasal epithelial populations, whereas ECC enriched for secretory and ciliated epithelial cell types. Cross-tissue comparison with publicly available healthy lung epithelial transcriptomics dataset revealed conserved mucociliary cell types alongside tissue-divergent transcriptional programs, including distinct regulation of ion transport and mucin-associated genes. Together, these findings demonstrate the importance of sampling modality in resolving cervical epithelial heterogeneity, and reveal tissue-specialized mucociliary regulatory programs underlying cervical mucus biology and fertility regulation.

**IN BRIEF:** The endocervix constitutes a critical mucosal interface within the female reproductive tract, where coordinated epithelial ion transport and mucus secretion regulate reproductive function while maintaining a selective barrier against ascending pathogens. Single-nucleus RNA sequencing (snRNA-seq) of healthy cervical tissues collected by Tischler punch biopsy and endocervical curettage enabled us to define the cellular and transcriptional organization of the healthy human cervix, revealing evidence of sampling-dependent epithelial heterogeneity and conserved mucociliary cell types with tissue-divergent transcriptional programs across the cervical and pulmonary mucociliary epithelium.

## INTRODUCTION

The human cervix is the lower, fibromuscular portion of the uterus that serves as a dynamic mucosal barrier between the uterine cavity and the vagina, regulating female reproductive tract (FRT) homeostasis, fertility, and pregnancy (Holdsworth-Carson et al., 2023; Monin et al., 2019). The cervix represents a critical site for both understanding reproductive pathology, such as sexually transmitted disease, dysplasia, and infertility, as well offers a potential target for pharmacologic intervention in the context of contraception (Han et al., 2017). The cervix is comprised of the endocervix, transitional zone, and an ectocervix, which are anatomically and histologically distinct regions (Ferenczy et al., 1987). The endocervix extends toward the uterine cavity and is lined by columnar epithelium, while the ectocervix projects into the vagina and consists of stratified squamous epithelium (Madhukaran et. al., 2025). Epithelial cells in the endo and ectocervix regulate interactions with the local microbiome by secreting mucins, maintaining barrier integrity, and expressing different levels of immunomodulatory factors (Tomasi et al., 2019). Unlike other mucin-producing organs such as the lung or intestine, cervical mucus is hormonally sensitive and the biophysical characteristics of the cervical mucus vary considerably as a function of cycle stage, age, and during pregnancy (Moghissi et. al., 1972; Najmabadi et al., 2021; Han et al., 2022). In addition to protecting the upper reproductive tract from ascending pathogens, cervical mucus plays a critical role in fertility by regulating sperm transport through the endocervix and provides a physiological marker of the fertile window (Lacroix et al., 2020). Disruption of cervical mucus function can contribute to cervical-factor infertility, which might account for an estimated 5-10% of female infertility cases (Poon et al., 1985).

Recent single-cell studies have shed light on the cellular, transcriptional, and regulatory shifts that occur in the cervix as a function of pregnancy and in the context of cancer in both human and mouse models. These studies have revealed distinct expression profiles of secretory cells, including changes in mucin-coding genes, genes related to barrier function, and overall shifts in the abundance of epithelial cell states across pregnancy in mice (Cooley et al., 2023). In the context of cancer, for example, studies have revealed diverse populations of cancer stem cells, as well as hallmarks of cervical cancers associated with HPV infection (Madhukaran et al., 2025). Recent efforts to standardize cell identities across the female reproductive tract have highlighted the need for comprehensive annotation of cervical cell populations to facilitate cross-study integration (Ong et al., 2026). Prior characterization studies of the cervix have also commonly used bulk RNA-sequencing methods that do not allow for a clear delineation of cellular and molecular signatures (Mukhopadhyay et. al., 2019; Pathare et. al., 2023). Finally, existing single-cell studies largely capture disease-associated states, including malignant transcriptional reprogramming, pathogen-driven immune activation, and organoid models with limited physiologically fidelity (Li et al., 2022; Peng et al., 2025; Prakash et al., 2025). However, to date, the transcriptional programs and the harmonized cell identities that define healthy cervical mucosal function remain incompletely characterized, thus limiting our understanding of the molecular mechanisms regulating cervical mucus production. Defining cellular and transcriptional programs in the context of healthy tissue is therefore a crucial step towards identifying molecular targets for advancing our understanding of the cervix and nominating interventional strategies that act locally within the cervix in disease, fertility, and pregnancy.

The anatomical accessibility of the cervix, particularly the ectocervix, allows direct sampling and also administration of a range of therapeutics, with a time-to-pharmacological response as little as two hours (Han et al., 2019). Cervical tissues are typically sampled using two methods. Small biopsies from the outer ectocervix or near the squamocolumnar junction can be obtained using Tischler forceps, or endocervical curettage (ECC) can be used for mechanical scraping of cells from the inner mucosal lining of the endocervical canal (Underwood et, al., 2012; Myriokefalitaki et. al., 2016; Massad et. al., 2023; Cooper et. al., 2023). These two sampling methods, by nature of where they sample and how they sample, will yield drastically different cell compositions and distinct immune profiles (Tomasi et al., 2016). Despite the widespread use of both sampling methods in clinical settings, primarily for cancer screening, differences between these two cervical sampling modalities have not been well-defined at single-cell resolution. This is important so that mechanistic studies with collected samples in both health and disease can have an informed baseline reference.

We fill this gap by creating a single-nuclei healthy human cervical transcriptomic reference map from both the endo and ectocervix (Parvez et al., 2026). Sampling method results in cell compositions that reflect established tissue biology, with direct implications for the interpretation of transcriptomic data derived from routine clinical biopsies. We identify distinct cell-cell communication networks that underpin each compartment, revealing differences in each epithelial sub-set. Finally, to identify features of the cervical mucosa that are distinct from other mucociliary epithelia, we integrated secretory and ciliated cells from our cervical atlas with publicly available lung mucociliary populations (Sikkema et. al., 2023). We used this integrated dataset to identify transcriptional programs conserved across both tissues and those specific to the cervix. Our data defines cells that can be obtained across two commonly used biopsy methods from the cervix and how these compare to the lung.

## MATERIALS AND METHODS

### Human Cervical Tissue Collection

Cervical tissue biopsies were obtained from four healthy, premenopausal female donors aged 28–45 years with a mean 34.5 ± 7.4 years (**Supplementary Table 1**). Participants met inclusion criteria if they were between 18–45 years of age regularly menstruating and not using any hormones (contraceptive or otherwise).

### Clinical Protocol for Cervical Tissue Biopsy Sample Collection

The cervix was visualized with a speculum and a scopette was used to wipe away any vaginal discharge. Each donor underwent dual-modality sampling using both an endocervical curettage (ECC) using a Kevorkian endocervical curette and punch biopsy using a Tischler biopsy forceps. Endocervical curettage was performed by inserting the curette approximately 1 cm into the endocervical canal and applying gentle circumferential abrasion around the full 360° cervical opening circumference. Tischler punch biopsy was subsequently performed by positioning the forceps at the squamocolumnar junction of the ectocervix and closing the instrument to excise the tissue specimen. All specimens were immediately snap-frozen via immersion in liquid nitrogen and stored at −80°C until nuclear isolation for snRNA-seq library preparation.

### Single-Nuclei Ribonucleic Acid Sequencing (snRNA-seq)

For single-nucleus RNA-seq, cells were lysed to release intact nuclei, with cytoplasmic contents subsequently removed, individual nuclei are then isolated and encapsulated for sequencing (Ding et al., 2020). Nuclei were isolated from eight frozen tissue samples using the Chromium Nuclei Isolation Kit (10x Genomics, PN-1000494) according to the manufacturer’s protocol (CG000505). Isolated nuclei were counted and assessed for integrity, then loaded onto Chromium Next GEM Chip On-Chip Multiplexing (OCM) chips with four samples multiplexed per channel using the Chromium GEM-X Single Cell 3ʹ OCM Reagent Kit. Single-nucleus libraries were prepared following the manufacturer’s protocol (CG000832) on the 10x Genomics Chromium X instrument. Libraries were quantified, pooled, and sequenced on an Illumina NextSeq 2000 platform, targeting a sequencing depth of approximately 30,000 read pairs per nucleus. Raw reads were aligned to the GRCh38 reference using Cell Ranger (Schneider et al., 2017). Filtered features–barcode matrices were imported into R and parsed into Seurat objects (R Core Team, 2026). Gene expression was restricted to protein-coding genes as defined by Ensembl GRCh38 release 111 (retrieved via biomaRt) to exclude non-informative non-coding features. Library preparation and sequencing were performed at the Genomics & Molecular Biology Shared Resource at the Dartmouth Cancer Center.

### Quality Control and Doublet Detection

Per-nuclei quality metrics were computed for each sample, including the number of detected genes, total UMI counts, and the percentage of reads mapping to mitochondrial and ribosomal genes. Adaptive thresholds were derived per sample using a median absolute deviation (MAD) framework (±5 MADs), with hard lower bounds of ≥ 200 genes and ≥ 500 UMIs per nuclei; nuclei exceeding the upper MAD boundary on any metric were excluded. Doublets were identified and removed with scDblFinder (v1.24.10), run on the SingleCellExperiment (v1.30.1) representation of each sample independently before merging.

### Dimensionality Reduction and Batch Correction

After filtering, counts were log-normalized and the top 3,000 highly variable genes were selected per sample by the variance-stabilizing transformation. Mitochondrial percentage, ribosomal percentage and total UMI count were regressed out during scaling. Principal Component Analysis (PCA) was followed by UMAP visualization on the top 30 PCs and Leiden clustering at resolution 0.5. This resolution was selected after evaluating multiple clustering resolutions and choosing the setting that provided stable separation of transcriptionally distinct cell populations. To correct for donor identity and sequencing run while deliberately preserving biopsy-type differences as a biological variable, three integration strategies were benchmarked in parallel: Canonical Correlation Analysis (CCA)-based integration and reciprocal PCA (rPCA) via Seurat (v5.5.1) and Harmony (v2.0.4). Integration quality was scored across biological conservation (silhouette, cLISI), donor batch correction and sequencing-run batch correction (iLISI, kBET, graph connectivity, principal-component regression), and the highest-scoring embedding was used for downstream analysis.

### Cell Type Annotation

Because a comprehensive reference for the cellular composition of the healthy human cervix remains limited, we manually curated a cell-type marker dictionary for annotation **(Supplementary Table 2)**. Ten populations were identified: Secretory Epithelial 1, Secretory Epithelial 2, Basal Epithelial, Suprabasal Epithelial, Ciliated Epithelial, Mesenchymal Stromal, Lymphatic Endothelial, Blood Vessel Endothelial, Myeloid, and Lymphoid cells. Cell identity was assigned using a three-tier annotation strategy comprising: (i) expert markers, genes selected by domain experts based on their known and observed cell-type specificity; (ii) canonical markers, defined as genes established in the literature as hallmarks of that cell type from mouse and human studies; and (iii) top marker genes identified from this dataset, defined as genes with the highest differential expression within each Leiden cluster. Integration of these three complementary sources of evidence enabled robust cell-type annotation based on established cervical and mucosal epithelial biology.

For orthogonal validation independent of the initial clustering, cell-type assignments were verified using Sargent (v1.0.1), a marker-based scoring algorithm that assigned cell identity based on the manually curated dictionary of cell-type specific marker genes (Nouri et al., 2023). Cell-type assignments from Sargent were compared with the initial cluster-based annotations using a confusion matrix and cell-type-specific Jaccard indices.

### Differential Expression Between Biopsy Sites

Differential expression between biopsy sites was assessed within each cell type by Wilcoxon rank-sum test (Seurat::FindMarkers (v5.5.1), ECC versus Tischler), restricted to genes expressed in at least 20% of cells of that cell type, which also defined the cell-type-specific expressed-gene background used for enrichment (Hao et. al., 2024). No fold-change pre-filter was applied during testing (logfc.threshold = 0); genes were called differential at adjusted *p* < 0.05 (Benjamini–Hochberg) and |log2 fold-change| > 0.25 (Benjamini et al., 1995). Cell types with fewer than 20 cells, or lacking both biopsy types, were not tested. For the differential-expression heatmap, whole-sample pseudobulk profiles were formed by summing raw counts per sample, restricted to the donors with matched Tischler and ECC biopsies, and variance-stabilized with DESeq2 (v1.51.3). Per-gene z-scores (clipped to ±2.5) of the top DEGs per cell type and direction were displayed; mitochondrial and ribosomal genes were excluded from this panel (Love et. al., 2014). Rows were ordered by cell-type block and direction and columns by biopsy site then donor, without hierarchical clustering.

### Pathway Over-Representation Analysis

Pathway enrichment for was performed using over-representation analysis (ORA). For each cell type, Gene Ontology Biological Process (GO:BP) enrichment was conducted separately for genes significantly upregulated in ECC and Tischler samples, defined as adjusted *p* < 0.05 and |log2 fold-change| > 0.25 (Ashburner et. al., 2000; The Gene Ontology Consortium, 2026). Enrichment was performed using clusterProfiler::enrichGO (v4.20.0) with Benjamini– Hochberg correction (Wu et. al., 2021). The background gene set consisted of genes detected in at least 20% of cells within the corresponding cell type, thereby restricting the analysis to genes eligible for differential-expression testing rather than all annotated genes. Canonical markers were identified independently using Seurat::FindMarkers (v5.5.1). The five most significant terms in each direction were displayed for each cell type.

### Inference of Cell–Cell Communication

Ligand–receptor signalling was inferred with CellChat (v2.2.0) using the integrated, normalized expression and cell-type labels and the CellChatDB.human database (Jin et. al., 2025). Over-expressed genes and interactions were identified, communication probabilities computed (triMean, with population-size correction), and cell-type populations contributing fewer than 10 cells removed by CellChat:filterCommunication. Pathway-level probabilities were aggregated and network centrality computed.

### Differential Enrichment of Curated Cervical Programs

To contrast biopsy sites, a panel of manually assembled gene sets spanning cervical epithelial identity, mucins and glycosylation, ion transport, hormone signalling, ECM and immune programs was tested within each epithelial subtype. For each subtype, log2 fold-changes between ECC and Tischler were computed for every expressed gene (Seurat::FoldChange; genes detected in ≥ 5% of either condition). For each gene set, the enrichment score was calculated as the mean log2 fold-change across detected genes. Gene sets with fewer than ten detected genes within a subtype were excluded. Statistical significance was assessed by comparing the distribution of gene-level log2 fold-changes within each gene set against all other expressed genes using a Wilcoxon rank-sum test, with Benjamini–Hochberg correction across gene sets within each subtype.

### Cross-Tissue Comparison with Lung Epithelium

Cervical epithelial nuclei were compared with a publicly available human lung snRNA-seq dataset (Sikkema et. al., 2023). The lung dataset was filtered to female donors, nuclear suspensions, and epithelial populations. Data were imported into R using reticulate (v1.46.0) and anndata (v0.8.0) and converted to a Seurat (v5.5.1) object. Ensembl gene identifiers were mapped to HGNC gene symbols using available feature annotations, with additional mapping performed using biomaRt (v2.66.2) and org.Hs.eg.db (v3.22.0) (Durinck et. al., 2009). Where multiple Ensembl identifiers mapped to the same gene symbol, the feature with the highest total UMI count was retained; original Ensembl identifiers were preserved in the feature metadata.

Lung epithelial annotations were mapped to the cervical epithelial classification scheme as detailed in **Supplementary Table 3**. The resulting framework comprised five cervical epithelial subtypes, four of which had corresponding populations in the lung; Secretory Epithelial 2 was retained as a cervix-specific population (Schupp et al., 2021). Cervical nuclei were restricted to these epithelial subtypes, and sampling modality was used as a proxy for cervical anatomical compartment, distinguishing Tischler punch biopsy from endocervical curettage (ECC) samples. Cervical and lung datasets were restricted to shared genes, merged, and integrated across tissues using canonical correlation analysis (CCA) implemented in Seurat (v5.5.1), with 3,000 anchor features and the first 30 principal components. Integration parameters (k.anchor and k.weight) were adjusted according to dataset size, and an unintegrated PCA representation was retained for comparison.

To assess transcriptional similarity across tissues, pseudobulk expression profiles were generated for each epithelial subtype and anatomical location, transformed to log-counts per million (logCPM), and compared within corresponding epithelial populations. Cross-tissue similarity was further quantified using MetaNeighbor (v1.29.0), with tissue specified as the study variable and similarity evaluated by area under the receiver operating characteristic curve (AUROC) (Crow et. al., 2018). For each shared epithelial subtype, each cervical sampling compartment was matched to the lung anatomical location with the highest within-subtype AUROC. Differential expression between each matched cervix–lung pair was then assessed using the Wilcoxon rank-sum test implemented in Seurat::FindMarkers, with genes required to be detected in at least 10% of cells (min.pct = 0.10). DE genes were defined by |log2 fold-change| > 0.25 and Benjamini–Hochberg-adjusted *p* < 0.05.

Gene Ontology Biological Process (GO:BP) over-representation analysis was performed separately on genes with higher expression in the cervix and lung for each matched comparison using clusterProfiler::enrichGO (v4.18.4). Enriched terms were summarized according to whether they were specific to the cervix, shared between tissues, or specific to the lung, and the top enriched processes were visualized for each matched comparison.

### Transcription-Factor (TF) Regulon Activity

TF regulon activity was estimated with VIPER (v1.46.0) on pseudobulk profiles aggregated by anatomical location as represented by the biopsy type × shared epithelial subtype (mean log-normalized expression) (Alvarez et. al., 2016). Regulons were taken from DoRothEA (dorothea_hs, confidence A/B) and converted with df2regulon; VIPER was run with method = “none” and a minimum regulon size of four (Garcia-Alonso et. al., 2019). The cross-tissue comparison used the four shared subtypes (Secretory Epithelial 1, Basal, Suprabasal and Ciliated Epithelial); Secretory Epithelial 2 is cervix-specific and was excluded. Tissue-divergent TFs were ranked by a divergence score equal to the absolute difference in mean NES between cervix and lung groups, up-weighted when activity changed sign across tissues, down-weighted when both tissues were concordantly positive or negative, and gated to require a maximum NES > 1; the top 50 TFs were retained and classified as cervix-enriched, lung-enriched or shared. Their DoRothEA A/B target genes belonging to curated ion-channel and mucin panels were assembled into a bipartite TF–target network (activation versus repression edges) target-gene expression was summarized as a dot plot across subtype × tissue groups.

### Statistical Analysis

Tests were two-sided unless stated otherwise, with Benjamini–Hochberg false-discovery-rate correction applied throughout. Significance is denoted as * adjusted *p* < 0.05, ** < 0.01, *** < 0.001; ns, not significant. A fixed seed (set.seed(42)) was used for reproducibility.

### Visualization

Figures were generated in R (v4.6.1) with ggplot2 (v4.0.3), patchwork (v 1.3.2), cowplot (v1.2.0), ComplexHeatmap (v2.28.0), circlize (v0.4.18), UpSetR (v1.4.0), ggVennDiagram (v1.5.7) and ggh4x (v0.3.1) (Wickham et al., 2016; Gu et al., 2016; Gu et al., 2014; Conway etal., 2017; Gao et al., 2021). UMAPs, dot plots, violin plots and marker heatmaps were produced with Seurat and ComplexHeatmap; chord diagrams with circlize; UpSet plots with UpSetR; and lollipop, tile-enrichment and bipartite-network panels with custom ggplot2 layers. Figures used a 9-pt base font (Helvetica/Arial) and were exported as vector PDF and 300-dpi PNG.

## RESULTS

### Single-Nuclei RNA-sequencing Defines the Cellular Composition of the Healthy Endocervix and Ectocervix

To map the cell types present in the healthy human cervix and to better understand how sampling method impacts cell populations retrieved, we performed snRNA-seq on paired Tischler and ECC biopsies across four healthy, premenopausal female donors (**Figure 1A**), resulting in a dataset comprised of 8,446 single nuclei. Due to insufficient data quality, one ECC sample was excluded from downstream analyses (**Supplementary Figure 1B**). A three-tiered cell-type annotation strategy (see **Methods** and **Supplementary Table 2**) was used to annotate cell types, and this resulted in ten transcriptionally distinct cell clusters including cells in the Epithelial lineage (Secretory Epithelial 1, Secretory Epithelial 2, Basal, Suprabasal and Ciliated), Stromal lineage (Mesenchymal Stromal), Vascular lineage (Lymphatic Endothelial, Blood Vessel Endothelial) and Immune lineage (Myeloid, Lymphoid) (Figure 1B and **C**). We recover all expected cell types that are well-defined by expected marker genes (Tomasi et al., 2019). Secretory Epithelial 1 cells were characterized by expression of the gel-forming mucins MUC5AC and MUC5B, whereas Secretory Epithelial 2 cells expressed SCGB1D2 and CFD. Basal Epithelial cells expressed TP63, KRT5, and COL17A1, while Suprabasal Epithelial cells were distinguished by KRT4, KRT13, and IVL. Ciliated Epithelial cells expressed FOXJ1 and motile cilia-associated genes, including TTC29 and DNAH12. Mesenchymal Stromal cells expressed PDGFRB, COL1A2, and ACTA2. Vascular populations comprised Lymphatic Endothelial cells expressing PROX1, CCL21, and LYVE1, and Blood Vessel Endothelial cells expressing PECAM1, PLVAP, and FLT1. Immune populations included Myeloid cells expressing CSF1R, CD68, and CYBB, and Lymphoid cells expressing CD3D, ZAP70, and GRAP2 (**Figure 1D**).

**Figure 1.**
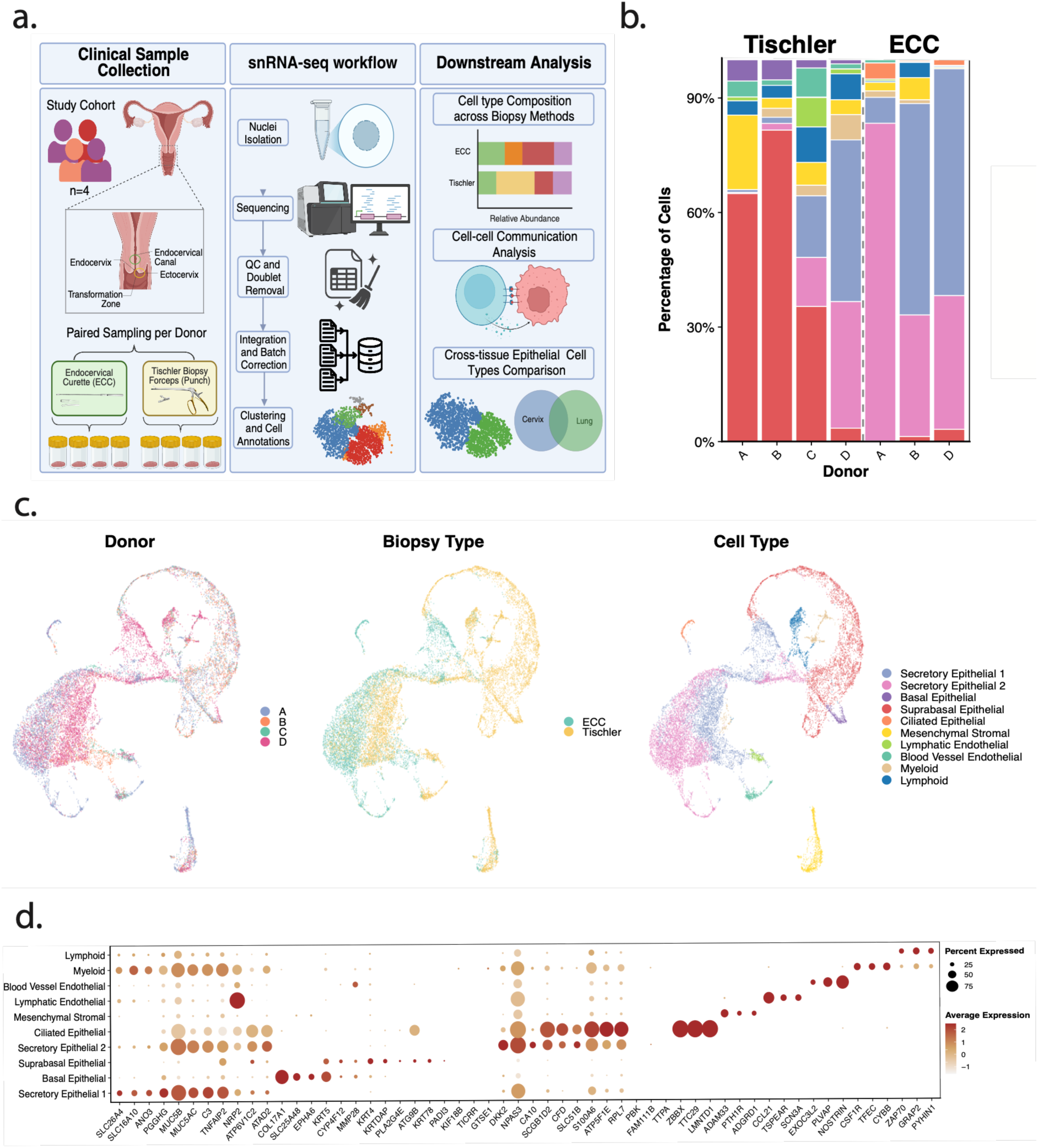
Single-nuclei transcriptomic profiling of the healthy human cervix sampled by Tischler punch biopsy and endocervical curette (ECC). (a) Overview of the study design, sample collection (Tischler from ectocervix & ECC from endocervix) and analysis workflow. Illustration created using BioRender.com. (b) Stacked bar charts showing the proportion of each annotated cell type per donor, stratified by biopsy site. Each bar represents one donor (A– D); cell types are color-coded as indicated in the legend. (c) Uniform Manifold Approximation and Projection (UMAP) of all cells after batch integration, coloured by donor identity (left), biopsy site (centre) and annotated cell type (right). (d) Dot plot showing scaled average expression (colour) and percentage of expressing cells (size) for marker genes identified using a 3-tiered cell type marker framework across the ten annotated cell types. Marker genes are grouped by cell type along the x-axis.

For orthogonal validation using Sargent, overall label concordance was 55.1%. Cells with ambiguous Sargent assignments, defined by low Gini indices, were classified as unassigned and manually inspected. The highest concordance was recorded in transcriptionally distinct populations (Ciliated Epithelial, Blood Vessel Endothelial, and Mesenchymal Stromal) and greater divergence among closely related epithelial subtypes. This independent, marker-based approach was concordant with our manually curated annotations for well-resolved cell types, while highlighting transitional or intermediate populations that require additional resolution. Taken together, we have constructed a high-quality reference dataset for the human cervix using two distinct biological sampling methods on the same donors.

We also found that cell type proportions varied substantially between the two sampling methods (**Figure 1B**), as expected based on anatomical location. The Tischler biopsy resulted in predominantly suprabasal, basal, and some secretory epithelial cell populations, while the ECC samples were predominantly enriched in secretory epithelial cells, including ciliated and mucin-secreting cell subtypes. Given the low abundance of lymphoid cells in our dataset, we are not powered to assess frequency differences in tissue resident CD8^+^ and CD4^+^ T cell compartments.

### Cervical Biopsy Method Defines the Transcriptional State of Cervical Mucinous Epithelium

To further investigate the extent of anatomical heterogeneity captured by the two cervical sampling modalities, we next assessed sampling-dependent transcriptional differences between ECC and Tischler samples within shared cell lineages. Secretory epithelial populations exhibited the greatest transcriptional divergence between sampling modalities, with Secretory Epithelial 1 and 2 showing 3,293 and 3,196 genes upregulated in ECC, compared with 2,302 and 1,401 in Tischler biopsies, respectively (**Figure 2A**). Myeloid cells similarly exhibited ECC-skewed differential expression, whereas suprabasal epithelial cells showed the opposite trend. This data suggests that the underlying transcriptomic differences between the endo and ectocervix are distinct across both epithelial and immune cell lineages.

**Figure 2.**
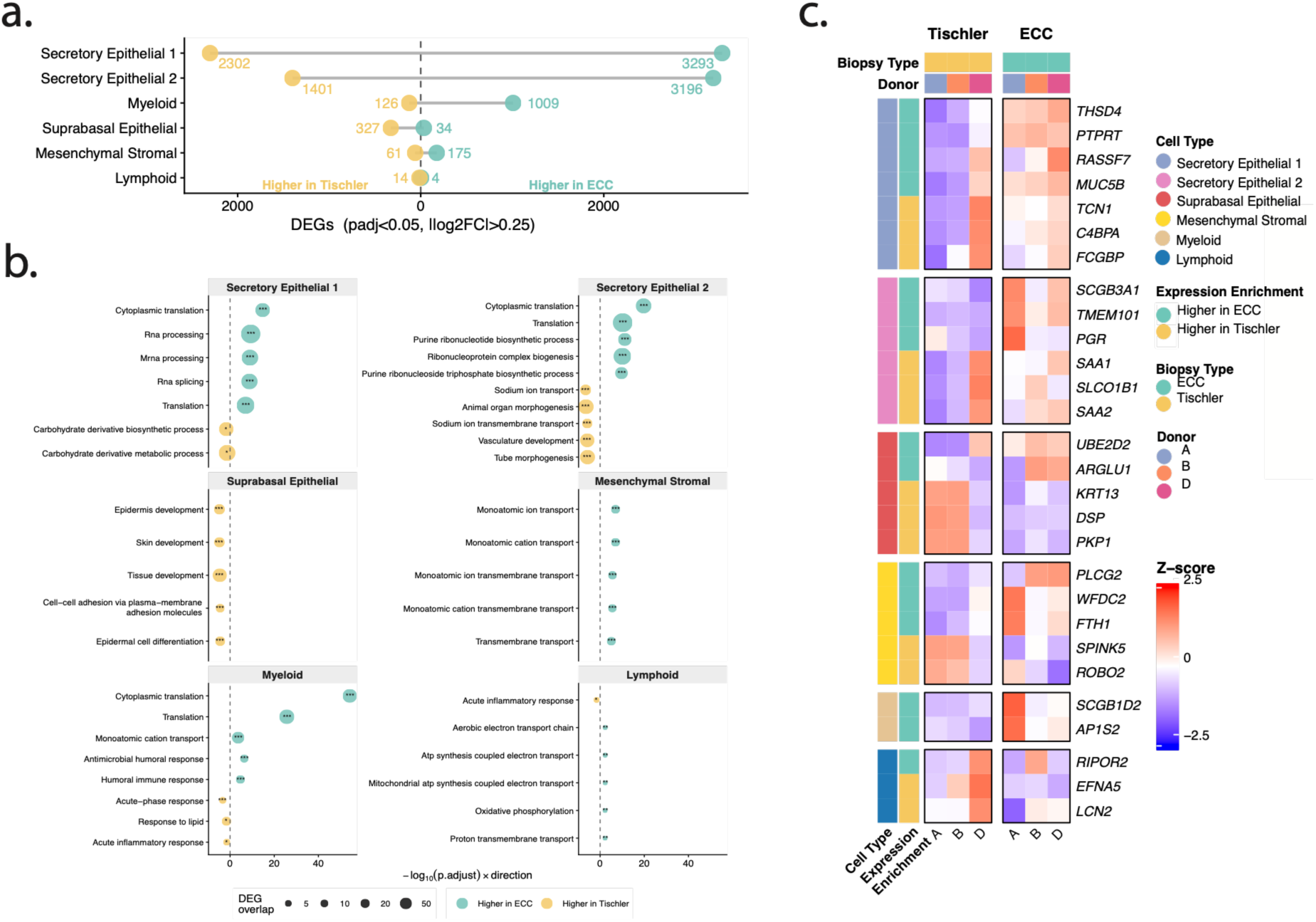
Transcriptional differences between Tischler and ECC biopsy sites across cervical cell types. **(a)** Lollipop plot showing the count of DEGs between biopsy sites in each cell type, identified by Wilcoxon rank-sum test (Seurat FindMarkers; ECC versus Tischler within each cell type; adjusted p < 0.05, |log2 fold-change| > 0.25) against a per-cell-type expressed-gene background. **(b)** Bubble plot of Gene Ontology Biological Process (GO) over-representation analysis (ORA) of ECC and Tischler upregulated genes, stratified by cell type. The x-axis represents signed −log₁₀(adjusted P) (positive, ECC; negative, Tischler), with dot size indicating DEG overlap and color denoting biopsy type. Enrichment used cell-type-specific gene backgrounds and pathway relevance filtering. Significance: adjusted P < 0.05, < 0.01, < 0.001. **(c)** Heatmap of donor-level pseudobulk expression for representative DEGs within each cell type. Raw counts were aggregated by donor and sampling modality, variance-stabilized using DESeq2, and standardized to gene-wise z-scores across samples. Rows are grouped by the cell type in which each gene was identified as differentially expressed and annotated according to DEG direction. Columns represent paired donor samples grouped by sampling modality.

Next, we wanted to determine if identified DEGs (p_adj_ < 0.01 and FDR < 0.01) were enriched for unique biological functions. We performed gene ontology over-representation analysis with each set of significant DEGs across each cell type (**Figure 2B**). We found that both secretory epithelial populations contained genes higher in ECC that were enriched for translation and RNA-processing-related terms like cytoplasmic translation, mRNA processing, RNA splicing, and ribonucleoprotein complex biogenesis. Tischler-derived secretory epithelial cells were enriched for carbohydrate derivative metabolism in secretory epithelial 1, and sodium ion transport, tube morphogenesis, and vasculature development in secretory epithelial 2 (**Figure 2B**). Suprabasal epithelial cells, which were located almost exclusively in Tischler-collected samples, were enriched for epidermal differentiation programs, including skin development and tissue development. Interestingly, we found that mesenchymal stromal cells from ECC samples were enriched almost entirely for ion transport processes. Finally, myeloid cells from ECC samples were mostly enriched for immune functions, including antimicrobial, humoral immune response terms, contrasting with an acute-phase/inflammatory response signature among the smaller set of Tischler-higher genes. The differences here may reflect the different immune functions of myeloid cells in the ectocervix as compared to the endocervix, where the former is characterized by tissue maintenance and the latter by response to the vaginal microbiome.

Next, we wanted to visualize DEGs that were significantly different across sampling method and drove pathway results for each cell type (**Figure 2C**). To do this, genes were selected from both ECC and Tischler-enriched DEGs sets (see **Methods**) and displayed as gene-wise z-scored expression across paired donor samples. Secretory epithelial populations showed the clearest separation by sampling modality. Several genes including THSD4, PTPRT, RASSF7, and MUC5B, show higher relative expression in ECC samples, whereas TCN1, C4BPA, SAA1 and FCGBP express highly in Tischler samples. These findings highlight considerable sampling-dependent transcriptional heterogeneity within shared cell lineages, potentially reflecting regional specialization across cervical compartments.

### Cell-Cell Communication Analysis Identifies Epithelial Subtype-Specific Signalling Networks

Given that we found distinct cell type differences across sampling method, and that each anatomical location performs distinct functions, we next wanted to leverage our data to understand how the communication networks differ (**Figure 3A**). Tischler biopsies exhibited broadly distributed signaling across epithelial, stromal, and immune populations, while ECC samples showed more concentrated interactions involving secretory epithelial and vascular cell populations consistent with their respective contributions in mucus secretion and barrier maintenance (Blaskewicz et al., 2011; Chumduri & Turco, 2021). Mean-ranked ligand-receptor pairs showed that pathways associated with epithelial homeostasis and immune surveillance accounted for the strongest interactions overall (**Figure 3B**), and incoming versus outgoing signaling for each epithelial subtype revealed that Secretory Epithelial 1 and 2 were predominantly signal-receiving populations, while Basal and Suprabasal Epithelial cells skewed toward outgoing signaling, suggesting a directional axis running from the basal/suprabasal compartment toward the secretory lineages (**Figure 3C**). We then dived deeper into understanding whether biopsy site shifted activity of curated cervical biology gene sets within each epithelial subtype (**Figure 3D**). Suprabasal Epithelial cells showed greater expression of squamous differentiation-associated programs in Tischler samples, while ECC samples exhibited elevated mucin and columnar epithelial signatures in secretory epithelial populations. These patterns are consistent with the differential anatomical access of Tischler biopsy and ECC to ectocervical and endocervical compartments, respectively.

**Figure 3.**
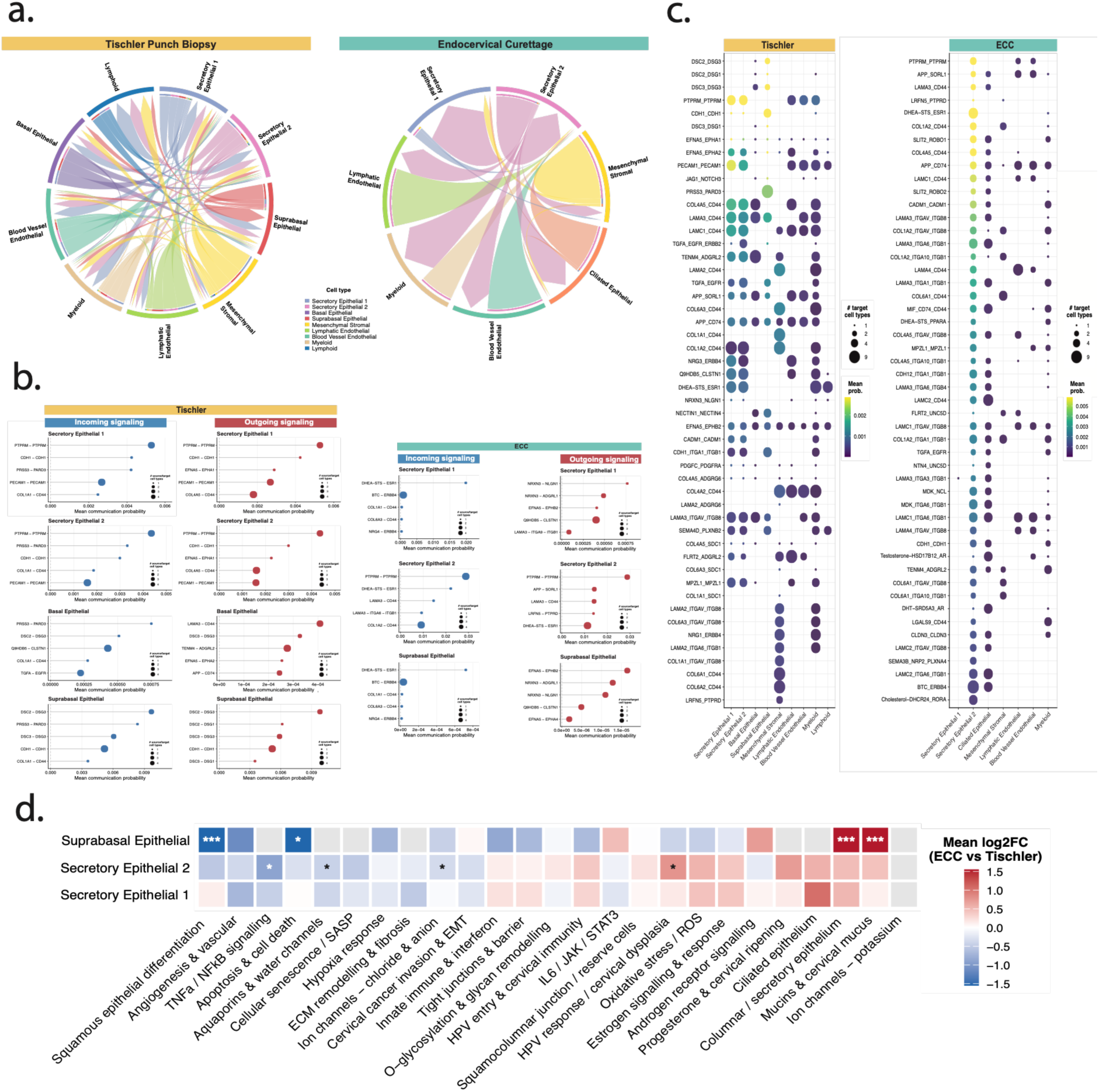
Cell–cell communication landscape and differential pathway programs in cervical epithelium. **(a)** Chord diagram summarizing aggregated cell–cell interactions inferred by CellChat across all annotated cell types. Ribbon and arc width are proportional to the aggregated interaction weight; cell types are color-coded as indicated in the legend. **(b)** Dot plot of the top predicted ligand–receptor interactions ranked by mean communication probability. Dot size indicates the number of target cell types involved; colour encodes mean communication probability. Interactions are ordered by overall strength. **(c)** Lollipop plots showing the top incoming (blue) and outgoing (red) ligand–receptor interactions, ranked by mean communication probability. Each point represents one ligand–receptor pair. **(d)** Differential pathway-enrichment heatmap contrasting ECC and Tischler biopsies within each epithelial subtype. For each curated cervical gene set, the enrichment score is the mean log2 fold-change (ECC versus Tischler) of its expressed genes (Seurat FoldChange over genes detected in ≥ 5% of either condition). Colour encodes direction and magnitude (red, higher in ECC; blue, higher in Tischler; grey, fewer than ten detected genes for that set in that subtype). Asterisks denote a competitive enrichment test (gene-set versus background log2 fold-changes, Wilcoxon rank-sum; Benjamini–Hochberg-adjusted: * < 0.05, ** < 0.01, *** < 0.001).

### Cervical Epithelial Subtypes Share Global Transcriptional Programs with Anatomically Distinct Lung Epithelium

The presence of mucociliary epithelia in the cervix and lung prompted us to ask whether these shared epithelial phenotypes have conserved transcriptional programs across tissues. To address this, we integrated our cervical nuclei dataset with a publicly available lung snRNA-seq dataset restricted to healthy female donors, mapping both organs onto a shared five-subtype epithelial dataset (**Figure 4A**). Overall, our dataset has good representation of each epithelial cell lineage across each sampling site (**Figure 4B**). Interestingly, we found that secretory epithelial 2 cells appear to be mostly identified in the cervix samples and basal epithelial cells to be present more abundantly in lung samples. This is likely due to how these tissues are sampled, but nonetheless, still enables a comparison across these cell states.

**Figure 4.**
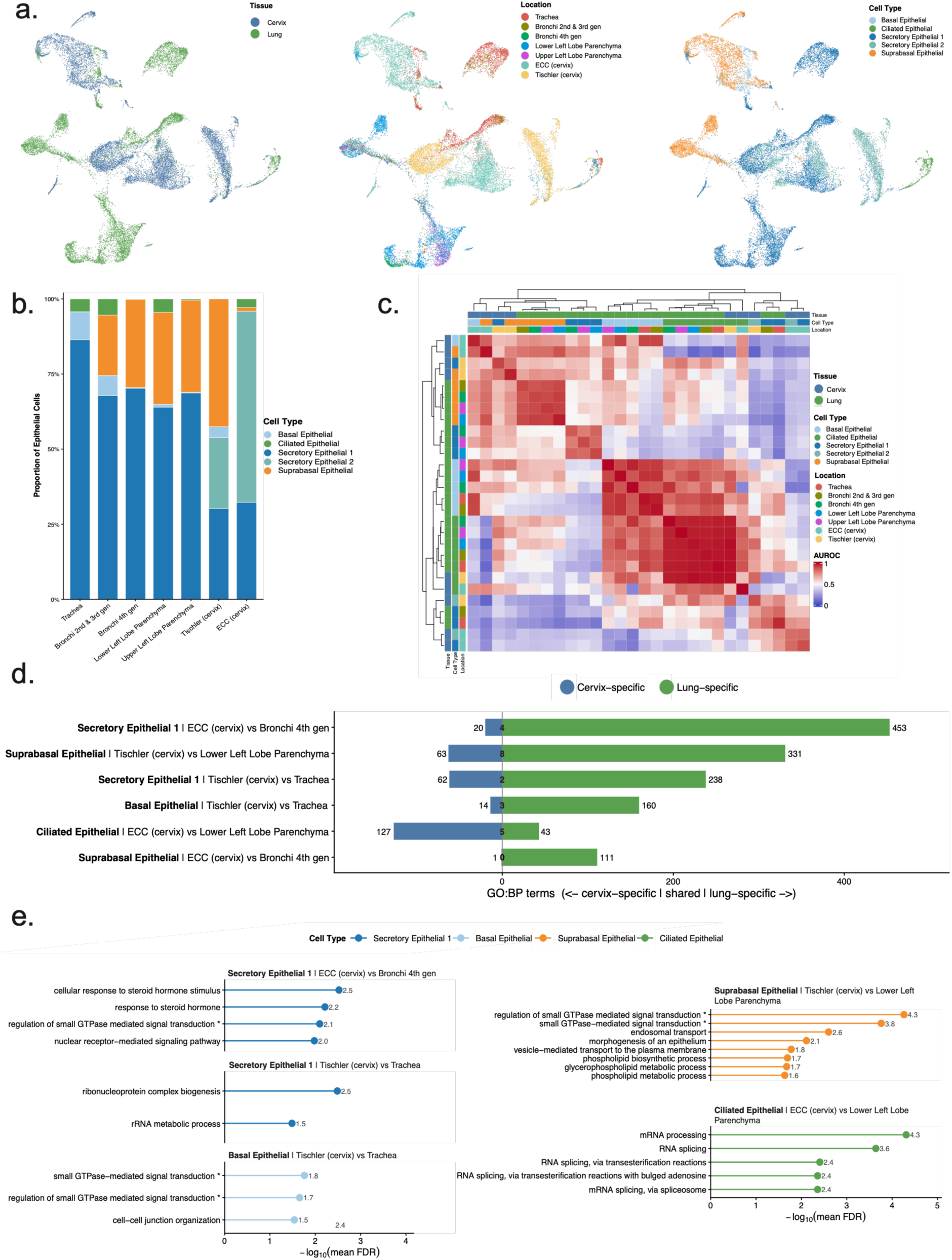
Cross-tissue analysis reveals shared and distinct Epithelial programs in the cervix and lung. **(a)** UMAP embeddings of CCA-integrated cervical and lung epithelial nuclei, coloured by tissue of origin (left), anatomical location (center) and shared epithelial subtype (right). Lung subtypes were mapped to the shared five-subtype scheme as described in Supplementary Table 1; each point represents a single nucleus. **(b)** Stacked bar chart of the proportional composition of shared epithelial subtypes across lung anatomical locations and cervix biopsy sites. **(c)** MetaNeighbor cross-tissue AUROC heatmap across pseudobulk groups defined by cell type and anatomical location, with tissue treated as the study label. AUROC ≈ 1 denotes populations that are highly reproducible across tissues. For each shared epithelial subtype, the maximum within-subtype AUROC identifies the lung anatomical location most similar to each cervix biopsy; these matched cervix–lung pairs are the contrasts analyzed in (d) and (e). **(d)** Diverging bar chart of Gene Ontology biological process (GO:BP) term counts for each AUROC-matched cervix-biopsy versus lung-location contrast (one row per contrast). The count of shared terms is shown at the center. Terms were identified by GO:BP over-representation analysis (enrichGO) of the genes higher in cervix and higher in the matched lung location for each pair (Wilcoxon rank-sum, |log2 fold-change| > 0.25, FDR < 0.05). **(e)** Top enriched GO:BP terms for representative matched contrasts, shown as lollipop plots of −log10(mean FDR). Each block corresponds to one cervix-biopsy versus lung-location comparison (annotated above the block) and points are colored by epithelial subtype. Asterisks mark terms passing the significance threshold.

Next, we performed a MetaNeighbor analysis to identify which populations of cells are most similar to each other across lung and cervix samples. We found that ciliated epithelial populations exhibited the strongest transcriptional similarity between the cervix and lung, while other epithelial subtypes displayed varied tissue-associated divergence (**Figure 4C**). Interestingly, clustering of our similarity matrix appeared to be driven more by cell type rather than location alone. However, Gene Ontology enrichment analysis of differentially expressed genes when comparing lung versus cervix populations revealed substantial functional divergence between matched cervical and pulmonary ciliated epithelial populations, with 127 cervix-specific, 43 lung-specific, and only five shared biological processes (**Figure 4D**). Shared biological processes were primarily associated with mRNA processing and RNA splicing (**Figure 4E**). We did find enriched hormone processes in Secretory epithelial 1 cells when compared to similar cells from the bronchi. Thus, despite conservation of epithelial identities on a global transcriptome level as assessed by transcriptome similarity, we found that epithelial cells of the cervix and lung adapt distinct transcriptional programs enriched for functions that mirror their respective mucosal environment.

### Transcription Factor Activity Reveals Tissue-Specific Ion Transport and Mucin Programs in Cervical Epithelium

Ion transport and mucin secretion are central to the hydration and biophysical properties of both cervical and airway mucus, prompting us to investigate whether their underlying transcriptional regulatory programs are conserved across tissues (Hill et. al., 2022; Han et. al., 2022). VIPER-inferred regulon activity analysis across the shared epithelial subtypes present in both tissues indicated that cervix and lung deploy a set of shared and different regulatory programs even within matched epithelial populations (**Figure 5A**). This includes cervix-enriched CCAAT/enhancer-binding protein delta (CEBPD) and lung-enriched heat shock transcription factor 1 (HSF1), peroxisome proliferator-activated receptor gamma (PPARG), transcription factor AP-2 alpha (TFAP2A), and melanocyte-inducing transcription factor (MITF). Additionally, several transcription factors, including GATA-binding protein 3 (GATA3), E2F transcription factor 1 (E2F1), PR/SET domain 14 (PRDM14), and androgen receptor (AR), form a highly active core regulatory block across both anatomical compartments and epithelial cell types.

**Figure 5.**
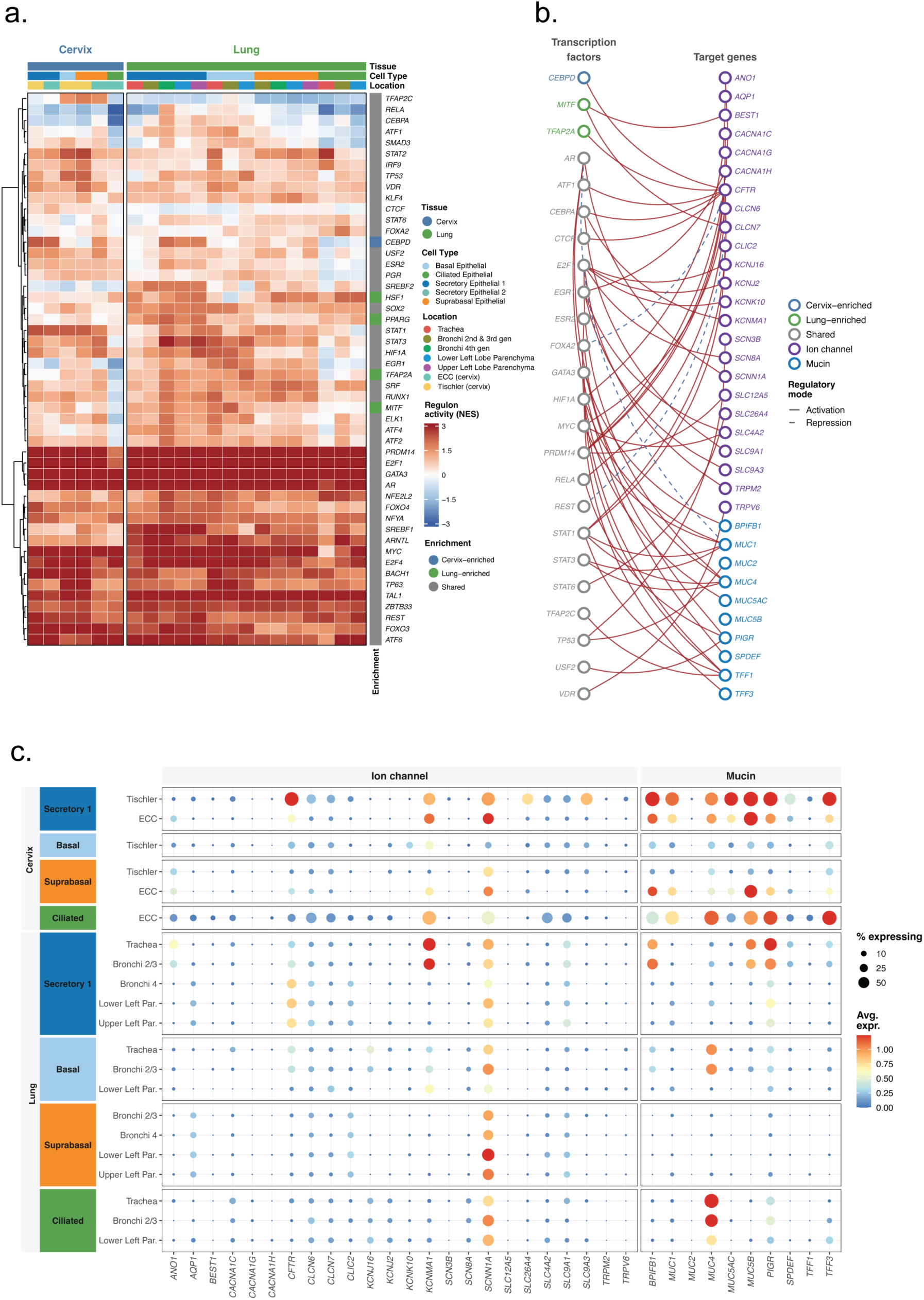
Tissue-divergent transcription factors regulate ion-channel and mucin gene programs in cervical and lung epithelium. **(a)** Heatmap of VIPER-estimated transcription factor (TF) regulon activity (normalized enrichment score, NES) across pseudobulk groups defined by anatomical location × shared epithelial subtype, for the four cross-tissue subtypes (Secretory Epithelial 1, Basal, Suprabasal and Ciliated Epithelial; Secretory Epithelial 2 is cervix-specific and excluded from the cross-tissue comparison). The top 50 TFs ranked by a cervix–lung divergence score are shown. Rows are hierarchically clustered (Ward’s method); columns are split by tissue and ordered by subtype and location. Regulons were taken from DoRothEA confidence levels A/B. Color scale spans NES −3 to +3. **(b)** Bipartite network diagram linking the tissue-divergent TFs in (a) to their ion-channel and mucin target genes, as defined by DoRothEA A/B regulons. **(c)** Dot plot showing average normalized expression (color) and percentage of expressing cells (size) for the TF target genes identified in (b) across the epithelial subtype × tissue groups.

We next wanted to map these transcription factors onto their ion-channel and mucin target genes in order to determine if any may play a role in mucus production, either in a shared or unique way across anatomical compartments. This analysis resulted in a bipartite regulatory network linking cervix-enriched, lung-enriched, and shared transcription factors to their downstream targets by predicted activating and repressive interactions (**Figure 5B**). We found several core transcription factors directly impact mucin genes, including several solute transporters and endomucins, such as mucin 5AC, oligomeric mucus/gel-forming (MUC5AC) and mucin 5B, oligomeric mucus/gel-forming (MUC5B). Interestingly, CCAAT enhancer binding protein delta (CEBPD), a cervix-enriched transcription factor, was directly linked to the ion channel CF transmembrane conductance regulator (CFTR). Target gene expression analysis further revealed distinct tissue- and subtype-dependent expression patterns (**Figure 5C**). In the cervix, CFTR and sodium channel epithelial 1 subunit alpha (SCNN1A) were most prominent within secretory epithelial populations, whereas potassium calcium-activated channel subfamily M alpha 1 (KCNMA1) was detected across multiple epithelial states. By contrast, anoctamin 1 (ANO1) and several other ion-transport genes showed relatively sparse expression. Mucin-associated genes similarly displayed pronounced subtype specificity, with secretory and ciliated cervical populations expressing distinct combinations of mucin 4, cell surface associated (MUC4), MUC5AC, MUC5B, polymeric immunoglobulin receptor (PIGR), and trefoil factor 3 (TFF3). Together, these findings suggest that shared mucociliary cell identities are associated with tissue-specialized transcriptional regulatory programs, and we nominate candidate regulators of cervical epithelial ion transport and mucin-associated functions for future investigation.

## DISCUSSION

The cervix and the production of cervical mucus is a central component of the female reproductive tract, yet the epithelial populations and local tissue contexts that shape its composition and function remain incompletely characterized, particularly in healthy women. By resolving cervical epithelial states at single-nuclei resolution and comparing samples obtained using distinct biopsy approaches, our study addresses this gap by defining how sampling strategy influences the cellular and transcriptional features recovered from the cervix. We further extend our work to ask how epithelial cells in the healthy cervix compare to healthy lung in order to better understand tissue-specific and shared factors that impact mucus production.

We establish defined benchmarks for what cell types are recovered by two routine methods for sampling the cervix. Physiologically, the ectocervix is predominantly lined by stratified squamous epithelium, whereas the endocervix is lined by a columnar epithelium containing secretory and ciliated cell populations (Chumduri & Turco, 2021; Deng et al., 2019). This difference is reflected in the choice of biopsy type as the Tischler biopsy forceps obtain targeted ectocervical biopsies through a punch-biopsy mechanism, producing tissue fragments that preserve local epithelial and stromal architecture. In contrast, ECC uses a fenestrated curette with sharpened edges to circumferentially scrape the mucosal surface of the endocervical canal, generating smaller, more fragmented tissue specimens enriched in endocervical glandular epithelium, including ciliated and mucin-secreting cell populations (Massad et al., 2023). Our results delineate the differences in cellular composition between biopsy and ECC specimens with snRNA-seq and we provide a clear demarcation of cell identities in these samples. This has direct practical consequences for the field to guide which biopsy type to use depending on the cell types that are required. For example, suprabasal epithelial cells were almost exclusively obtained from Tischler biopsies, representing both the unique location and sample type. Additionally, by focusing on healthy women, our study helps establish a baseline cellular framework for understanding the cervix as a mucosal surface in its normal physiological state, against which disease-associated transcriptional changes can be more accurately interpreted. Our data also suggests that biopsy method should be clearly described and treated as covariate in a cervical single-cell study design if multiple techniques are used.

Using our dataset, we then compare cells across the two biopsy types and to another anatomical location, the lung. Our cell–cell interaction analysis on cervical tissues revealed distinct communication patterns among cervical epithelial populations, with basal and suprabasal cells contributing to inferred outgoing ligand–receptor signaling and secretory populations showing predicted incoming signaling. These findings are consistent with emerging single-cell studies of the cervical epithelium, in which basal and parabasal populations have also been identified as major sources of inferred signaling through pathways including BMP, TGF-β, MIF, and semaphorin signaling (Huang et. al., 2025). Our analysis extends existing evidence that cervical epithelial subtypes occupy distinct signaling roles, while further suggesting that secretory populations may represent important recipients of epithelial-derived signals in the healthy cervix, particularly the endocervix. Interestingly, the decrease in signaling complexity when comparing the ecto to the endocervix may reflect the fundamental difference in functions. The ectocervix, sampled with the Tischler biopsy method, is facing the vagina and exposed to the vaginal microbiome. The signaling architecture that coordinates proper immune defence and barrier function may therefore be much more complex as compared to the endocervix, as our data suggests (Madhukaran et al., 2025; Tomasi et al., 2019). However, since these interactions are inferred from ligand and receptor expression, they should be interpreted as candidate communication relationships rather than direct evidence of signaling and will require spatial and functional validation for confirmation. However, our analyses position the secretory and suprabasal compartments as dominant hubs of the cervical microenvironment and suggests a model in which structural epithelial layers actively regulate the differentiated secretory and ciliated populations responsible for mucus production and transport, rather than acting as passive scaffolding. This also suggests that hormonal sensitivity may itself be mediated through both direct steroid receptor and indirect (paracrine) fashion depending on the cell-type (Buchanan et al., 1998; Chung et al., 2013). Notably, the suprabasal transcriptional state was discinct across biopsy sites, indicating that structural epithelial compartments are dynamic components of the cervical mucosal environment. This interpretation is consistent with studies demonstrating that cervical epithelial homeostasis and differentiation are led by spatially organized epithelial lineages and local signaling programs, including Wnt, Notch, TGF-β, and EGFR signaling (Chumduri et al., 2021; Zhang et al., 2021). Links between CCIs, cervicovaginal mucus production and the composition of the vaginal microbiome should be more comprehensively explored in the future, particularly in the context of inflammation in the ectocervix (Zierden et al., 2023).

Finally, through a cross-tissue comparison, we found that core aspects of secretory and ciliated epithelial identities are either unique or conserved between the cervix and lung, positioning the cervix within a broader framework of mucosal epithelial biology. This finding is consistent with emerging cross-tissue single-cell atlases showing that epithelial cell types can retain conserved molecular programs across anatomically distinct organs while simultaneously acquiring tissue-specific transcriptional states (Elmentaite et al., 2022; Tabula Sapiens Consortium, 2022). In particular, multi-ciliated cells exhibit highly conserved ciliary gene-expression programs across respiratory and reproductive epithelia, suggesting that specialized epithelial functions can converge on common molecular architectures despite distinct tissue origins (Chae et. al., 2023). Our findings extend this to the cervix and demonstrate that cervical secretory and ciliated epithelial cells share core features with their airway counterparts while retaining a small set of cervix-specific transcriptional factor programs. These findings also corroborate existing proteomic studies of the mucus secretome where key secreted products, in particular gel forming mucins MUC5B and MUC5A are shared (Han et al., 2021). We also found that, despite global transcriptome similarity and a shared core of transcription factors, the cervix and lung exhibited distinct transcriptional regulatory programs. This may explain the distinct character of cervical mucus, which is more viscous than lung mucus, and nominates a smaller set of cervix-specific transcription factors, like CEBPA, that may regulate ion channels and mucins in the cervix (Rapp et al., 2024; Roberts et al., 2023). CEBPA has been documented to play a role in uterine corpus endometrial carcinoma, one of the most common tumors of the reproductive system, but its role in healthy cervix biology remains unexplored (Wang et al., 2024). Future work should pursue functional validation of these tissue-specific candidate regulators and extend this resource to characterize steroid influenced changes both within and between tissues so that the pathways of hormonal sensitivity in mucosal tissues can be better characterized.

This study has several limitations that must be considered when interpreting the findings. It is worth noting that the cohort size (n=4) is limited and while paired within-donor sampling strengthens the biopsy-method comparison, the donor pool was not balanced across menstrual cycle phase (3 luteal vs 1 follicular), limiting our ability to fully disentangle cycle-stage effects from biopsy-method effects on the transcriptional landscape. Second, this is a cross-sectional dataset collected at a single time point per donor so it does not capture temporal variation across the menstrual cycle, which would be necessary to establish how stable these cell-type-specific transcriptional programs are longitudinally (Burns et al., 2026; Wang et al., 2020). Third, our cross-tissue comparison relied on a publicly available lung dataset generated in a separate cohort and sequencing platform. Despite batch-aware integration and AUROC-based matching, residual cross-study technical variation cannot be fully excluded as a contributor to the tissue-divergent signatures we observed.

In summary, our findings establish that the healthy cervical epithelial populations combine conserved features of mucociliary tissues with tissue-specific programs adapted to reproductive function. This study provides a foundation for future work in cervical physiology and nominates both tissue-specific and shared regulators of mucus production by defining how clinical sampling shapes the cervical cell populations observed and by separating cross-organ conservation from cervical-specific regulatory programs. This work should help bridge molecular regulation of the cervical epithelium to the physical properties of mucus that govern barrier function, sperm transport and, ultimately, enable opportunities for locally acting therapeutics.

## DATA AVAILABILITY

The single-nucleus RNA sequencing data generated in this study have been deposited in the Gene Expression Omnibus (GEO) under accession number GSE349708, available at https://www.ncbi.nlm.nih.gov/geo/query/acc.cgi?acc=GSE349708.

## CODE AVAILABILITY

Analyses were performed in R (v4.6.1; R Core Team, 2026). The packages used are listed in **Supplementary Table 4**. The complete analysis pipeline, including preprocessing, clustering, and cell-type annotation code, is available at https://github.com/Goods-Lab/healthy-human-cervix-snRNAseq under an MIT license.

## ETHICS DECLARATION

The authors declare no conflict of interest.

## ACKNOWLEDGEMENTS

The authors acknowledge the Genomics and Molecular Biology Shared Resource at Dartmouth Cancer Center for providing genomic services, including the 10x Genomics Chromium single-cell GEM-based assay. Additionally, during the preparation of this work, Anthropic’s Claude and OpenAI’s ChatGPT was used to assist with minor copyediting and improve the clarity of select sentences. All AI-assisted text was critically reviewed and revised by the authors, who take full responsibility for the accuracy, originality, and scientific integrity of the manuscript. Generative AI was not used to generate experimental data, conduct computational analyses, or independently formulate scientific conclusions.

## AUTHOR CONTRIBUTIONS

A.P. contributed to data generation, data curation, formal analysis and writing (original draft). O.K. contributed to data curation, formal analysis, writing (review & editing). L.L. and R.L. contributed to project administration, clinical sample handling, and experimental design. T.C.T.L. contributed to validation. L.L.H. and B.A.G. acquired funding and provided supervision for writing (review & editing) and analyses. L.L.H. conceived of the study. All authors approved the final manuscript for submission.

