## Supplemental Files for "Tissue-specific mucociliary regulatory programs in the healthy human endocervix and ectocervix"

*Equal Contribution
[+]Co-Corresponding

**CO-CORRESPONDENCE**

**Supplementary Table 1. Donor demographics and hormonal profiling.** All donors underwent paired biopsy collection using both Tischler & ECC modalities at a single visit. Serum estradiol (E2) and progesterone (P4) levels were measured on the day of biopsy collection. Menstrual cycle phase was estimated using cycle day and serum P4 concentrations, with P4 ≥ 3 ng/mL used as a study-defined criterion supporting luteal-phase assignment, consistent with clinical guidance that elevated progesterone provides presumptive evidence of recent ovulation (ASRM Practice Committee, 2021).

| Donor ID | Age (years) | Cycle Day | Cycle Phase | E2 (pg/mL) | P4 (ng/mL) |
|---|---|---|---|---|---|
| **A** | 45 | 11 | Follicular | 219 | 0.83 |
| **B** | 34 | 18 | Luteal | 133 | 19.31 |
| **C** | 28 | 25 | Luteal | 176 | 21.27 |
| **D** | 31 | 19 | Luteal | 133 | 8.10 |

**Supplementary Table 2. Three-tiered gene-marker based cell-type annotation framework for the human cervix.** Cell-type identities were assigned using three complementary sources of marker-gene evidence: (i) **expert markers**, genes selected by domain experts based on their known and observed cell-type specificity; (ii) **canonical markers**, established cell-type markers supported by published literature from mouse and human studies; and (iii) **key marker genes**, highly differentially expressed genes identified within individual Leiden clusters identified in the single-nucleus RNA-sequencing (snRNA-seq) dataset. Gene symbols follow standard nomenclature.

| Cell Type | (i) Expert Markers | (ii) Canonical Markers | (iii) Key Marker Genes |
|---|---|---|---|
| Secretory Epithelial 1 | MUC5AC, MUC5B, KRT7 | MUC5AC (Gipson et al., 1997), MUC5B (Gipson et al., 2001), MUC16 (Andersch-Bjorkman et al., 2007), SPDEF (Park et al., 2007), LCN2 (Friedl et al., 1999) | SLC26A4, SLC16A10, ANO3, PGGHG, C3, TNFAIP2, ATP6V1C2, ATAD2 |
| Secretory Epithelial 2 | SCGB1D2, CFD, S100A6 | SCGB1D2 (Zafrankas et al., 2006), SCGB2A1 (Becker et al., 1998), KRT7 (Senzaki et al., 1997), KRT19 (Senzaki et al., 1997) | CA10, CFD, SLC51B, S100A6, ATP5F1E, RPL7 |

| | | | |
|---|---|---|---|
| Basal Epithelial | TP63, COL17A1, KRT15 | KRT5 (Lloyd et al., 1995), KRT14 (Lloyd et al., 1995), TP63 (Yang et al., 1999), COL17A1 (McGrath et al., 1995), ITGA6 (Stepp et al., 1990), KRT15 (Waseem et al., 1999) | COL17A1, SLC25A48, EPHA6, KRT5, CYP4F12, MMP28 |
| Suprabasal Epithelial | KRT13, KRT4, IVL | KRT13 (Malecha et al., 1992), KRT4 (Leube et al., 1988), IVL (Sassoon et al., 1985) | KRT4, KRTDAP, PLA2G4E, ATG9B, KRT78, PADI3 |
| Ciliated Epithelial | KRT8, CFAP43, DNAH12, TEKT1 | FOXJ1 (You et al., 2004), DNAH12 (Yang et al., 2025), TEKT1 (Ryan et al., 2018), CFAP43 (Tang et al., 2017) | ZBBX, TTC29, LMNTD1, KIF18B, TICRR, GTSE1, PBK |
| Mesenchymal Stromal | DCN, COL1A2, COL3A1, LUM, PDGFRB | COL1A2 (Kinchen et al., 2018), ACTA2 (Skalli et al., 1986), MYLK (Yin et al., 2006), PDGFRB (Hellstrom et al., 1999), COL3A1 (Kinchen et al., 2018), LUM (Vorstandlechner et al., 2020), DCN (Pulkkinen et al., 1990) | MEG3, PDGFRB, COL3A1, COL6A3, DKK2, NPAS3 |
| Lymphatic Endothelial | PROX1, FLT4/VEGFR3, PDPN/Podoplanin, LYVE1 | PROX1 (Wigle et al., 1999), FLT4/VEGFR3 (Kaipainen et al., 1995), PDPN/Podoplanin (Breiteneder-Geleff et al., 1999), LYVE1 (Banerji et al., 1999), NRP2/Neuropilin-2 (Yuan et al., 2002), CLDN5/Claudin-5 (Baluk et al., 2007) | NRP2, PROX1, FLT4/VEGFR3, PDPN/Podoplanin, LYVE1, CCL21, TSPEAR, SCN3A |
| Blood Vessel Endothelial | FLT1/VEGFR1, PECAM1/CD31 | PECAM1/CD31 (Muller et al., 1989), FLT1/VEGFR1 (Kaipainen et al., 1993) | ADAM33, PTH1R, ADGRD1, EXOC3L2, PLVAP, NOSTRIN, FLT1/VEGFR1, RAMP3 |
| Myeloid | CD163, CD68, FCN1 | CD163 (Kristiansen et al., 2001), CD68 (Holness et al., 1993), CD1C (Dzionek et al., 2000), LYZ (Miyauchi et al., 1985), FCN1 (Lu et al., 1996) | MYO1F, CSF1R, TFEC, CYBB |
| Lymphoid | CD3D, CCR7, NCAM1/CD56 | CD3D (Gold et al., 1987), CCR7 (Sallusto et al., 1999), CD4 (Rogozinski et al., 1984), NCAM1/CD56 (Lanier et al., 1989), | ZAP70, GRAP2, PYHIN1, PTPRC, FYB1, LCP2 |

**Supplementary Table 3 | Mapping of lung epithelial annotations to the shared cervix–lung subtype scheme.** Lung cells were first restricted to include only epithelial nuclei from female donors (sex = female, suspension_type = nucleus, Celltypes_master_high = Epithelia). Fine annotations (obs column Celltypes_master_higher_immune) were then mapped to the five-subtype scheme shared with the cervix. Secretory Epithelial 2 is cervix-specific and has no lung counterpart. Non-epithelial contaminants were excluded from downstream analyses.

| Lung Annotation | Mapped Cervical Subtype | Rationale |
|---|---|---|
| Epi_Basal | Basal Epithelial | Basal keratinocyte-like airway progenitors. |
| Epi_secretory | Secretory Epithelial 1 | Airway secretory cells. |
| Submucosal_Glands | Secretory Epithelial 1 | Mucous/serous secretory gland cells. |

| | | |
|---|---|---|
| AT2 | Secretory Epithelial 1 | Surfactant-secreting alveolar type 2 cells. |
| Ciliated | Ciliated Epithelial | Multiciliated cells. |
| AT1 | Suprabasal Epithelial | Alveolar type 1 cells with flattened morphology. |
| Secretory Epithelial 2 | No Lung Mapping | Cervix-specific subtype; excluded from cross-tissue comparison. |

**Supplementary Table 4 | R Packages Utilized.** Summary of R packages and their corresponding versions used for single-nucleus RNA sequencing data processing, statistical analysis, and visualization.

| Package | Role | Version |
|---|---|---|
| R | Base language/environment | 4.5.2 |
| Seurat/SeuratObject | snRNA-seq processing, DE (Wilcoxon), FoldChange | 5.5.1 |
| harmony | Integration benchmarking | 2.0.4 |
| scDblFinder | Doublet detection | 1.24.10 |
| SingleCellExperiment/scran/scuttle | QC/SCE handling | 1.32.0/1.38.1/1.20.0 |
| DESeq2 | Pseudobulk VST (Fig 2c), variance stabilisation | 1.51.3 |
| clusterProfiler/org.Hs.eg.db/AnnotationDbi | GO:BP ORA (enrichGO), gene mapping | 4.18.4/3.22.0/1.72.0 |
| CellChat | Cell–cell communication (Fig 3a–c) | 2.2.0 |
| dorothea/viper | TF regulons / regulon activity (Fig 5) | 1.22.0/1.44.0 |
| MetaNeighbor/SummarizedExperiment | Cross-tissue AUROC matching (Fig 4) | 1.29.0/1.40.0 |
| reticulate/anndata/ rhdf5 | Lung .h5ad import | 1.46.0/0.8.0/ 2.54.1 |
| biomaRt | Protein-coding filter, Ensembl↔symbol | 2.66.2 |
| fgsea/msigdbr | Gene-set resources (legacy/optional) | 1.36.2/26.1.0 |
| ComplexHeatmap/ circlize | Heatmaps, chord diagrams | 2.26.1/0.4.18 |

| | | |
|---|---|---|
| UpSetR/ggVennDiagram | UpSet / Venn diagrams | 1.4.0/1.5.7 |
| ggplot2/patchwork/ cowplot/ggh4x | Plotting / composition | 4.0.3/1.3.2/1.2.0/0.3.1 |
| dplyr/tidyr/tibble/ stringr/forcats | Data wrangling | tidyverse 2.0.0 |
| Matrix/matrixStats/future | Sparse algebra/stats /parallel | 1.7.5/1.5.0/1.70.0 |
| svglite/ragg/scales/ RColorBrewer/viridis | Devices/scales/palettes | 2.2.2/1.5.2/1.4.0/1.1.3/0.6.5 |

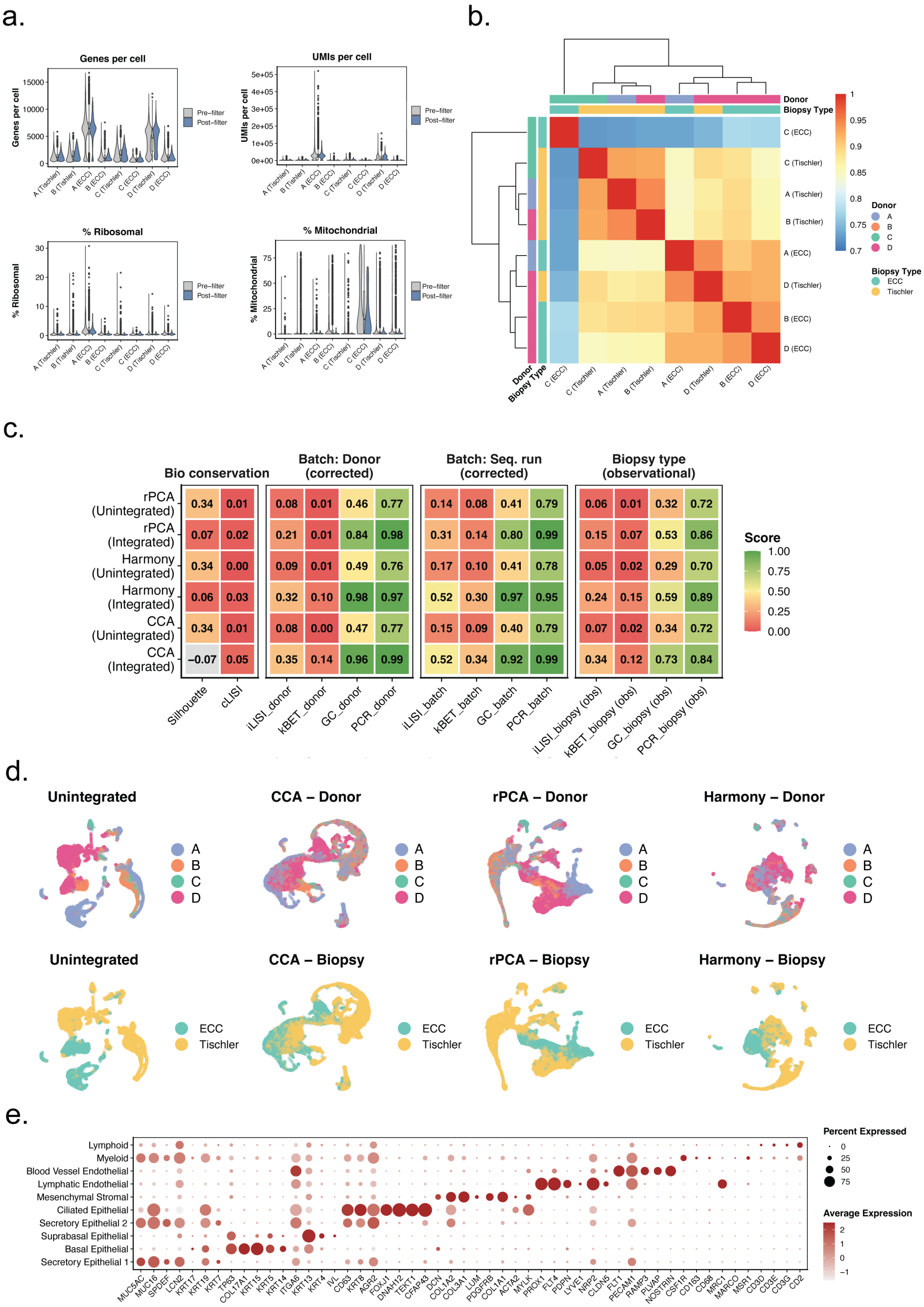
a.
Genes per cell
UMIs per cell
% Ribosomal
% Mitochondrial
Pre−filter
Post−filter
A (Tischler)
B (Tishler)
A (ECC)
B (ECC)
C (Tischler)
C (ECC)
D (Tischler)
D (ECC)
b.
Donor
Biopsy Type
C (ECC)
C (Tischler)
A (Tischler)
B (Tischler)
A (ECC)
D (Tischler)
B (ECC)
D (ECC)
ECC
Tischler
c.
Bio conservation
Batch: Donor (corrected)
Batch: Seq. run (corrected)
Biopsy type (observational)
rPCA (Unintegrated) 0.34 0.01 0.08 0.01 0.46 0.77 0.14 0.08 0.41 0.79 0.06 0.01 0.32 0.72
rPCA (Integrated) 0.07 0.02 0.21 0.01 0.84 0.98 0.31 0.14 0.80 0.99 0.15 0.07 0.53 0.86
Harmony (Unintegrated) 0.34 0.00 0.09 0.01 0.49 0.76 0.17 0.10 0.41 0.78 0.05 0.02 0.29 0.70
Harmony (Integrated) 0.06 0.03 0.32 0.10 0.98 0.97 0.52 0.30 0.97 0.95 0.24 0.15 0.59 0.89
CCA (Unintegrated) 0.34 0.01 0.08 0.00 0.47 0.77 0.15 0.09 0.40 0.79 0.07 0.02 0.34 0.72
CCA (Integrated) −0.07 0.05 0.35 0.14 0.96 0.99 0.52 0.34 0.92 0.99 0.34 0.12 0.73 0.84
Silhouette
cLISI
iLISI_donor
kBET_donor
GC_donor
PCR_donor
iLISI_batch
kBET_batch
GC_batch
PCR_batch
iLISI_biopsy (obs)
kBET_biopsy (obs)
GC_biopsy (obs)
PCR_biopsy (obs)
Score
d.
Unintegrated
CCA – Donor
rPCA – Donor
Harmony – Donor
CCA – Biopsy
rPCA – Biopsy
Harmony – Biopsy
e.
Lymphoid
Myeloid
Blood Vessel Endothelial
Lymphatic Endothelial
Mesenchymal Stromal
Ciliated Epithelial
Secretory Epithelial 2
Suprabasal Epithelial
Basal Epithelial
Secretory Epithelial 1
MUC5AC MUC16 SPDEF LCN2 KRT17 KRT19 KRT7 TP63 COL17A1 KRT15 KRT5 KRT14 ITGA6 KRT13 KRT4 IVL CD63 KRT8 AGR2 FOXJ1 DNAH12 TEKT1 CFAP43 DCN COL1A2 COL3A1 LUM PDGFRB COL1A1 ACTA2 MYLK PROX1 FLT4 PDPN LYVE1 NRP2 CLDN5 FLT1 PECAM1 RAMP3 PLVAP NOSTRIN CSF1R CD163 CD68 MRC1 MARCO MSR1 CD3D CD3E CD3G CD2
Percent Expressed
Average Expression

**Supplementary Figure 1 | Quality control, batch correction benchmarking, and cell type annotation of the cervical epithelium samples.**

**(a)** Violin plots showing four quality control metrics (genes per cell, UMIs per cell, percentage ribosomal reads, and percentage mitochondrial reads) before (light grey) and after (blue) filtering, for each donor–biopsy sample. Filtering thresholds were applied uniformly across all samples.

**(b)** Hierarchically clustered heatmap of pairwise Pearson correlations between pseudobulk gene expression profiles for each donor–biopsy sample. Annotation bars indicate donor identity and biopsy type.

**(c)** Integration benchmarking table comparing three integration methods (rPCA, Harmony, CCA) in unintegrated and integrated states across four evaluation criteria: biological conservation (silhouette coefficient, cLISI), batch correction for donor (iLISI, kBET, graph connectivity, PCR), batch correction for sequencing run (same metrics), and biopsy-type mixing as an observational measure. Scores are color-coded from red (low) to green (high).

**(d)** UMAP embeddings comparing unintegrated data and three integration methods (CCA, rPCA, Harmony), colored by donor identity (top row) and biopsy site (bottom row), illustrating the degree of batch correction achieved by each approach.

**(e)** Dot plot showing scaled average expression (color) and percentage of expressing cells (size) for curated cell type marker genes used to annotate all clusters. Markers are grouped by cell type along the x-axis; rows represent annotated cell types.

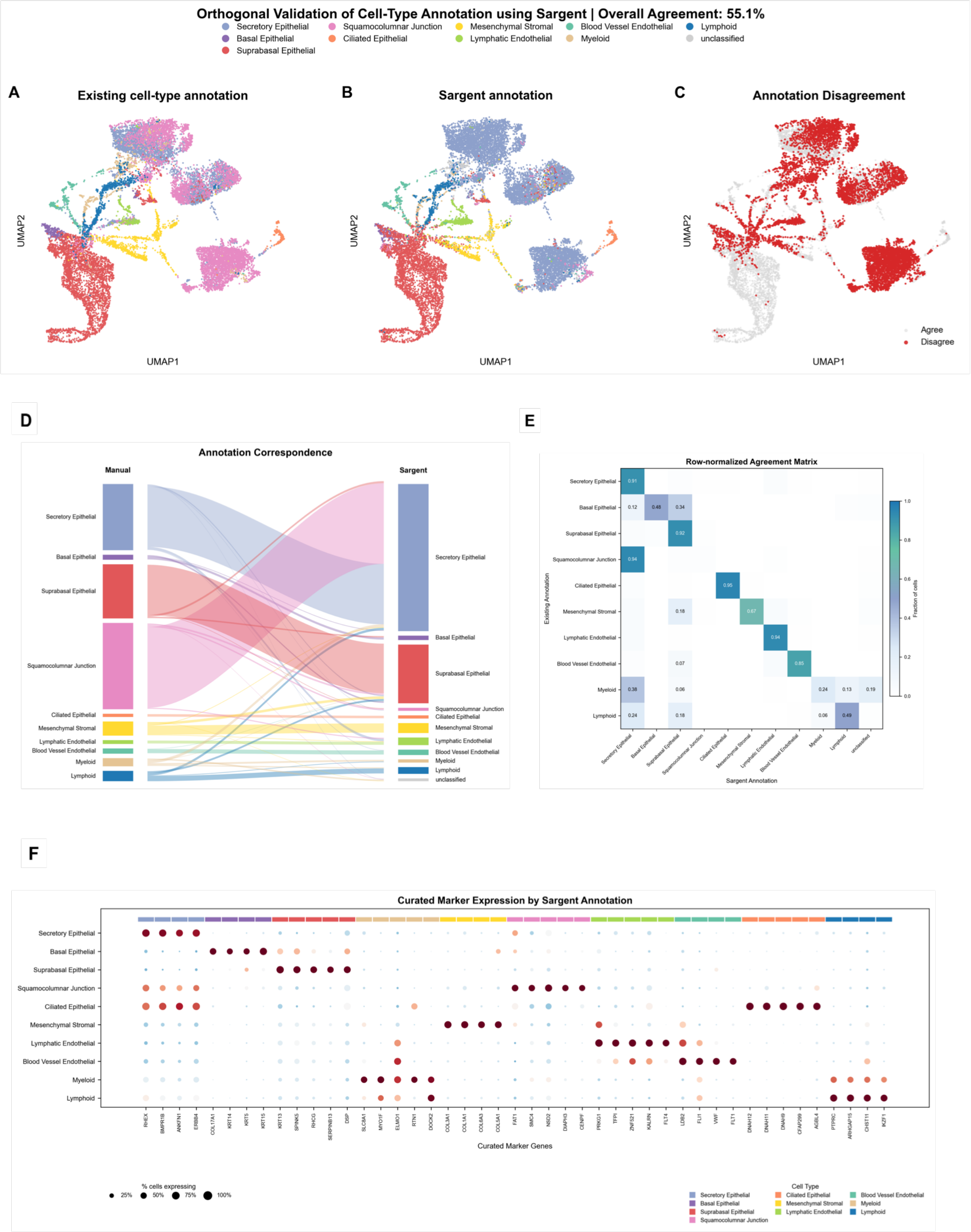


**Supplementary Figure 2 Orthogonal validation of manual cell-type annotation using Sargent.** (A) UMAP embedding colored by existing (manual) cell-type annotation. (B) UMAP embedding colored by Sargent-derived cell-type annotation. (C) UMAP embedding highlighting cells with agreement (gray) versus disagreement (red) between manual and Sargent annotations; overall agreement was 55.1%. (D) Sankey diagram showing correspondence between manual and Sargent cell-type assignments. (E) Row-normalized agreement matrix showing the fraction of cells from each manually annotated cell type assigned to each Sargent-predicted cell type. (F) Dot plot of curated marker gene expression across Sargent-annotated cell types, showing percentage of cells expressing each marker (dot size) grouped by cell type (color bar).
